# Pharmacologic inhibition of Protein Kinase C α and Protein Kinase C β halts renal function decline indirectly, by blunting hyperphagia, and directly reduces adiposity in the ZSF1 rat model of type 2 diabetes

**DOI:** 10.1101/2020.12.02.406892

**Authors:** Ju Wang, Agustin Casimiro-Garcia, Bryce G. Johnson, Jennifer Duffen, Michael Cain, Leigh Savary, Stephen Wang, Prashant Nambiar, Matthew Lech, Shanrong Zhao, Li Xi, Yutian Zhan, Jennifer Olson, James A. Stejskal, Hank Lin, Baohong Zhang, Robert V. Martinez, Katherine Masek-Hammerman, Franklin J. Schlerman, Ken Dower

## Abstract

Type 2 diabetes (T2D) and its complications can have debilitating, sometimes fatal consequences. Despite advances that address some of the metabolic aspects of T2D, for many patients these approaches do not sufficiently control the disease. As a result, an emerging therapeutic strategy is to target the pathobiological mechanisms downstream of T2D metabolic derangement that can result in organ damage, morbidity, and mortality in afflicted individuals. One such proposed mechanism involves the Protein Kinase C (PKC) family members PKCα and PKCβ, which have been linked to diabetes-induced tissue damage to organs including the kidneys. To evaluate the therapeutic potential of dual inhibition of PKCα and PKCβ in the context of T2D, we have evaluated a potent and orally bioavailable inhibitor, herein referred to as Cmpd 1, in the ZSF1 rat model of leptin-receptor deficiency, obesity-driven T2D. Therapeutic dosing of Cmpd 1 virtually halted renal function decline but did so indirectly by blunting the hyperphagia response of these animals. Beyond this clear but indirect effect, Cmpd 1 had direct and prominent effects on body weight and in liver and inguinal white adipose tissue (iWAT) when administered to ZSF1 obese rats.

## Introduction

Diabetes affects an estimated 170 million people worldwide, posing an enormous humanitarian and economic challenge^1–3^. Current estimates are that complications from diabetes account for 1 in every 8 dollars spent on medical care in the United States^1^. The major cause of morbidity and mortality in diabetic individuals stems from hyperglycemia-induced macro- and microvascular damage that leads to cardiovascular disease, damage to organs such as eyes and kidneys, and damage to neurons. Diabetes can be difficult to control. As a result, large numbers of individuals continue to succumb to diabetes-induced tissue damage that can result in limb amputations, blindness, end-stage renal disease (ESRD), and death. Indeed, the Centers of Disease Control estimates that diabetes is the 7^th^ leading cause of death in the United States, with recent reports indicating that this may be an underestimation^4^. A clear and present need exists for therapies that limit the extent of tissue and organ damage that occurs in the diabetic state^2,5^.

A particularly serious complication of diabetes is diabetic nephropathy (DN), which is characterized by cellular and structural aberrations in the kidney. These include hypertrophy and expansion of mesangial cells, deposition of extracellular matrix, thickening of the glomerular basement membrane, and effacement of podocytes^6^. DN is the leading cause of ESRD in the United States and is also a major risk factor for heart attack and stroke^7,8^. Several signaling molecules and pathways have been implicated in DN progression (reviewed in^8,9^). Amongst these are the Protein Kinase C (PKC) family of serine-threonine kinases. PKCs consists of conventional family members α (alpha), β (beta, with β1 and β2 splice variants), and γ (gamma); novel family members δ (delta), ε (epsilon), θ (theta) and η (eta); and atypical family members ζ (zeta) and ι/λ (iota in humans, also referred to as lambda in rodents). Conventional and novel PKC family members are activated directly by diacylglycerol (DAG), the levels of which increase in diabetes due to *de novo* synthesis^10,11^. They are also indirectly activated by oxidative stress and advanced glycosylation end products (AGEs), which increase during diabetic tissue injury^12,13^. Because the activation of PKCs themselves can result in the production of reactive oxygen species (ROS), it is postulated that PKC activation in the diabetic setting can lead to a destructive feed-forward loop that causes cell and tissue damage^14^.

At this time, the strongest link between PKCs and DN progression is with the conventional PKCs PKCα and PKCβ, which have been implicated to human and preclinical DN through elevated expression (PKC α and β) and genetic association (PKCβ)^15–22^. Studies using single and double knockout mice in models of streptozotocin (STZ)-induced type 1 diabetes (T1D) indicate non-redundant roles for these two PKCs in promoting DN: namely, PKCα in the loss of podocyte integrity and increased permeability of the glomerular filtration barrier; and PKCβ in renal cell hypertrophy and fibrosis^23–28^. Furthermore, administration of staurosporine-based PKC inhibitors Ruboxistaurin or CGP41251 is reno-protective in rodent models of T1D (STZ) and/or T2D (db/db model)^16,26,29^. Importantly, however, data suggest that some PKC family members may play protective roles in the kidney. For example, mice deficient in the novel PKC PKCε develop spontaneous kidney disease^30^, and the atypical PKCs PKCs ζ and ι/λ are required for the maintenance of proper podocyte polarity and function^31,32^. Therefore, pharmacologic inhibition of PKCα and PKCβ, one that effectively spares other PKC family members such as PKCε and the atypical PKCs, could constitute a promising therapeutic strategy for DN.

To investigate this, we evaluated an orally bioavailable and selective PKCα/β dual inhibitor, Cmpd 1^33^, in the ZSF1 rat model of T2D. ZSF1 rats are obtained from a cross of two strains heterozygous for leptin receptor mutations^34^. F1 progeny that are not homozygous for leptin receptor deficiency are normophagic. These animals do not develop disease, and although technically not lean are referred to as ZSF1 lean rats. F1 progeny that are homozygous for leptin receptor deficiency are hyperphagic, and become obese on high carbohydrate diet. ZSF1 obese rats develop metabolic syndrome by 8 weeks of age, proteinuria by 12 weeks of age, and renal disease by 24 weeks of age^34–40^. ZSF1 obese rats are one of relatively few rodent models of T2D characterized by DN progression to ESRD, with death resulting at 45-50 weeks of age, and are being increasingly used as a translationally relevant model to evaluate pharmacological agents^41–44^. Here we report our findings of therapeutic dosing of Cmpd 1 in the ZSF1 obese rat model.

## Materials and Methods

### Animals

ZSF1 (ZSF1-Lepr^fa^Lepr^cp^/Crl) lean rats (strain code #379) and ZSF1 obese rats (strain code #378) were obtained from Charles River Laboratories (Wilmington, MA). Chow containing Cmpd 1 was formulated by Research Diets Inc. (New Brunswick, NJ) by mixing 0.744 g Cmpd 1 per kg chow (Purina 5008 chow; 26.5% kcal from protein, 17.0% kcal from fat, 56.5% kcal from carbohydrates), followed by irradiation. All procedures involving animals were reviewed and approved by the Pfizer Institutional Animal Care and Use Committee.

### In vitro *kinase profiling*

Half maximal inhibitory concentrations of Cmpd 1 were determined by Z’-Lyte biochemical assays (Life Technologies SelectScreen^™^ Kinase Profiling Services, Thermo Fisher Scientific). Kinome selectivity was determined at 1 μM Cmpd 1 concentration against a panel of 119 kinases.

### In vitro *assays*

Glomeruli were isolated using an established protocol of serially sieving minced kidney cortex tissue through mesh filters of decreasing pore size^45^. For radioactive pan-PKC assays, glomeruli were lysed in a buffer consisting of 50 mM Tris-HCl pH 8, 5 mM EDTA, 5 mM EGTA, protease and phosphatase inhibitor cocktails, 1% 2-mercaptoethanol, and 0.028% n-dodecyl-β-D-maltopyranoside (Anatrace D310) by sonication using two 10 second pulses with a probe sonicator. Supernatants were collected following centrifugation at 4°C at 20,200 x g for 10 minutes. A 25 μL aliquot was used in a 50 μL ^32^P-ATP pan-PKC radioactive kinase assay with the pan-PKC pseudo-substrate peptide RFARKGSLRQKNV. The reaction was incubated for 10 minutes at 37°C prior to filter binding and scintillation counting. Data are reported as CPM incorporation per μg of total protein in the reaction, as determined using Pierce Coomassie Plus Protein Assay of the original lysate (Pierce 23236). Where indicated, the reaction was run in the presence of increasing amounts of Cmpd 1.

For the phorbol 12-myristate 13-acetate (PMA)-luminol assay, glomeruli isolated from PBS-perfused kidneys were diluted in an HBSS solution containing 1 mM luminol, 0.1 M HEPES, and 0.1% fatty-acid free BSA. Approximately 3,000 glomeruli were plated per well in a 96-well white OPTI-plate (Perkin Elmer 6005290). The reaction was started by the addition of PMA to a final concentration of 16 μM in the absence or presence of increasing Cmpd 1. Luminol luminescence was read on an Envision plate reader in kinetic read mode.

For the HEK293 cell-based assay, HEK293 cells engineered to over-express full-length PKCβ2 were stimulated with 3 nM PMA in DMEM media supplemented with 0.5% FBS. 4-hour cell culture supernatants were analyzed for IL-8 by MSD (Rockville, MD).

### Pharmacokinetic (PK) and food consumption studies

Three male 12-week old ZSF1 obese rats were fed control or 0.744g/kg Cmpd 1 chow for 4 days. Food consumption was estimated by measuring food provided versus food remaining in the hopper. Blood was collected through tail vein from each animal at 9 a.m. and 4 p.m. two days after chow dosing started, and at 9 a.m. four days after chow dosing started, for PK determination. Plasma protein binding of Cmpd 1 was measured by equilibrium dialysis of the *in vitro* fraction unbound of Cmpd 1 at a concentration of 2 mM in plasma from pooled male and female CD-1 mice. Plasma was centrifuged and supernatant was analyzed by liquid chromatography–tandem mass spectrometry to detect Cmpd 1 (York Bioanalytical Solutions; York, U.K.).

To evaluate food consumption in different rat strains, cohorts of 10 male ZSF1 obese rats, 8 male ZSF1 lean littermate rats, and 10 age-matched male Sprague Dawley (SD) rats were divided into two groups and fed chow with or without 0.744g/kg Cmpd 1 for 7 days.

### 10-week efficacy study

Male ZSF1 obese rats that had been maintained on Purina 5008 chow were switched to Purina 5008 chow with or without 0.744g/kg Cmpd 1 at 22 weeks of age. The study duration was 10 weeks. Estimated food consumption monitored weekly. An additional group of ZSF1 obese rats was included which received an amount of Purina 5008 chow to match the previous week’s estimated food consumption by the Cmpd 1 group. Five male ZSF1 lean littermates served as baseline controls. Intermediate blood samples were taken through tail vein. 24-hour urine samples at weeks 0, 5 and 10 were collected from animals in metabolic cages. Animals were sacrificed by CO2 asphyxiation followed by blood collection via cardiac puncture and tissues fixation in 10% neutral buffered formalin for histology. Unfixed inguinal white adipose tissue was also collected at study termination for downstream molecular analyses.

### Serum cholesterol, HDL, LDL, triglyceride, glucose, and insulin assays

All assays except the insulin assay were run with the ADVIA 1800 Chemistry System (Siemens Medical Systems Diagnostics, Tarrytown, NY, USA). The serum insulin assay was run using a SpectraMax® Plus384 Absorbance Microplate Reader (Molecular Devices San Jose, CA, USA). Siemens Medical System assay kits 10376501, 10311891, 10335892, and 10335891 were used for cholesterol, low density lipoprotein (LDL)-cholesterol, triglycerides, and glucose, respectively. Roche HDL-C Reagent 3rd generation assay kit 4713257190 was used for high density lipoprotein (HDL)-cholesterol assay. For insulin, an ALPCO Diagnostics assay kit 80-INSMS-E01 was used.

### Histopathology

Samples of formalin-fixed, paraffin embedded (FFPE) liver and kidney were sectioned and stained for Hematoxylin and Eosin (HE) and evaluated microscopically by a board-certified veterinary pathologist blinded to the treatment groups. In the kidney, microscopic features were assigned an end stage renal disease (ESRD) severity score of 1-5 (minimal to severe) using the following criteria: Grade 1 (minimal): < 10% of parenchyma affected, few glomeruli affected and no Bowman’s capsule thickening with only few protein casts and basophilic tubules observed; Grade 2 (mild): 10-25% of renal parenchyma affected, there is scattered thickening of Bowman’s capsule, +/- synechiation; Grade 3 (moderate): 26-50% of renal parenchyma affected, glomerular and tubulointerstitial changes are present; Grade 4 (marked): 51-75% of renal parenchyma affected, glomerular and tubulointerstitial changes are present; Grade 5 (severe): > 75% of renal parenchyma affected, glomerular and tubulointerstitial changes are present. In the liver, hepatocellular vacuolation was assessed on HE-stained sections and assigned a severity score of 0-3 (no macrovacuolation to moderate macrovacuolation). FFPE inguinal white adipose tissue was sectioned and stained for HE and the pan-leukocyte marker CD45 was detected by anti-CD45 ab10558 (Abcam, Cambridge MA; 1:5000 dilution) and red chromogen on the Leica Bond Auto-stainer Bond-Rx using heat induced epitope retrieval 2 - pH 9 (HIER 2) for 20 minutes. Positive controls consisted of banked FFPE rat lymph node and GALT. Whole-slide images of the immuno-stained sections were obtained with a Leica AT2 whole slide scanner (Leica Microsystems GmbH) for use in image analysis. CD45 immuno-positive areas were quantified using Definiens Tissue Studio image analysis software (Definiens AG, Germany) and quantitative data were reported as percentage of CD45 IHC stain vs total tissue section areas. For adipocyte size measurements, whole-slide images of the HE stained sections were obtained with a Leica AT2 scanner (Leica Microsystems GmbH) and adipocyte size was assessed by measuring cross sectional area (CSA) of adipocytes using newCAST software (Visiopharm, Broomfield, CO). Briefly, 10% meander sampling of the whole slide images were selected, point probe was used to identify adipocytes to be measured and nucleator probe was used to measure the CSA of the adipocytes. Data were reported as mean CSA of adipocytes in the section from each animal. HE and CD45-stained slides and quantitative image analysis data were analyzed and interpreted by a board-certified veterinary pathologist.

### RNA sequencing of inguinal white adipose tissue

Total RNA from inguinal white adipose tissue was isolated using RNeasy Plus Universal Kits (Qiagen 73404). Rat genome and Ensemble gene annotations were downloaded from Ensembl^46,47^. The clean raw sequence reads in FASTQ format were analyzed using the QuickRNASeq pipeline^48^, and reads were mapped to the rat reference genome (rn6) using STAR v2.4.0h^49^. Uniquely mapped reads were counted towards individual genes using featureCounts^50^. Genes that had no sequence reads mapped to them in 50% of samples were labeled as no or low expressed, and thus omitted from the differential expression analysis. This filtering step was included to reduce the number of false positives in the differential analysis^51^. A gene count table was generated by featureCounts and the differential analysis was performed by R packages edgeR 3.10.2^52,53^and Limma/voom 3.22.10^54^. Genes with a fold change greater than 1.5 or less than -1.5 and adjusted p-value (Benjamini-Hochberg)^55^ less than 0.05 were reported as significantly differentially expressed genes (DEGs).

### Pathway analysis

Pathway analysis was performed using QIAGEN Ingenuity Pathway Analysis (IPA). Two distinct Core Analyses were run: Obese versus Lean, and Cmpd 1 versus Food matched. A p-value less than 0.05 was set as the significance threshold for enriched pathways. A z-score algorithm was applied to determine if an enriched pathway was up- or down-regulated based on the input DEGs. A further Comparison Analysis between the two Core Analyses was run, and pathway enrichment findings were filtered for opposing trend and z-score in Obese versus Lean and Cmpd 1 versus Food matched. The Canonical Pathways and Diseases and Functions reported are those with a non-zero z-score in both comparisons, and p-value less than 0.05 and absolute z-score of 2, in at least one Core Analysis. The reported causal network regulators were all those with network-bias corrected p-value less than 0.05 in either Core Analysis.

### Statistics

Where shown, all error bars designate minus/plus Standard Deviation. For all analyses excluding RNA sequencing analyses (see above), p-values were calculated using 1-way or 2-way Anova with Bonferroni’s multiple comparison test.

## Results

### Cmpd 1 is a potent, ATP-competitive inhibitor of PKCα and PKCβ

Cmpd 1, the structure of which is not disclosed here, is an example of a 3-Aminopyrrolo[3,4-c]pyrazole-5(1H,4H,6H)carboxaldehyde ATP-competitive inhibitor^33^. **Figure 1A** provides a summary of the half maximal inhibitory concentration (IC_50_) of Cmpd 1 in enzymatic assays against PKC family proteins. Cmpd 1 inhibited PKCα and PKCβ with IC_50_ values of 2 and 11 nanomolar (nM), respectively. Cmpd 1 had moderate activity against some other PKC family members, notably PKCθ (IC_50_ 28 nM) and PKCγ (IC_50_ 96 nM). Selectivity against all other PKC family members was greater than 40-fold. Kinome profiling revealed good overall selectivity for Cmpd 1 against a panel of 119 kinases (**Figure 1B**). In a cell-based assay using HEK293 cells over-expressing PKCβ2, Cmpd 1 inhibited the production of IL-8 in response to the PKC activator phorbol 12-myristate 13-acetate (PMA) with an IC_50_ value of 26 nM (**Figure 1C**). In summary, Cmpd 1 is a potent, cell-permeable PKCα/β dual inhibitor with reasonably good selectivity over other PKC family members and the kinome overall.

**Figure 1.**
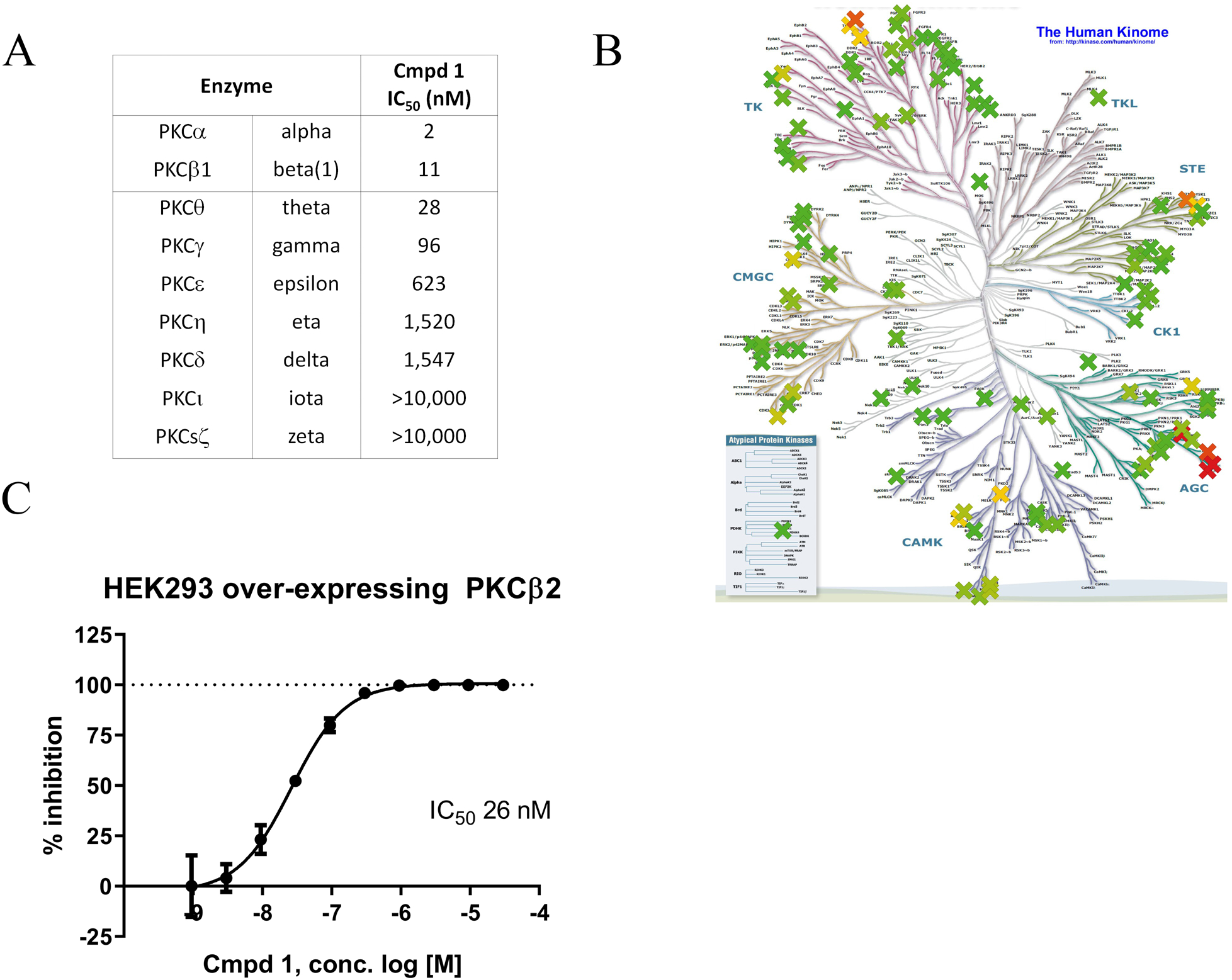
Compound 1 is a potent PKCα/ß dual inhibitor with selectivity over other PKC family members and the kinome overall. (A) Half maximal inhibitory concentration (IC_50_) of Cmpd 1 against PKC family members in enzymatic assays. (B) Kinome tree selectivity profiling of Cmpd 1 against a panel of 119 kinases. The percent inhibition by Cmpd 1 at 1 μM testing concentration is color coded, and ranges from 0% inhibition (darker green) to 100% inhibition (darker red) with 50% inhibition displayed in yellow. (C) Percent inhibition by Cmpd 1 of PMA-induced IL-8 in HEK293 cells overexpressing PKCß2.

### Cmpd 1 inhibits PKC activity in assays using glomeruli as source material

The roles of PKCα and PKCβ in DN have been attributed to functions in podocytes and mesangial cells, respectively, within the kidney glomerulus^23–28^. Based on our previously published transcriptomics analysis^40^, all PKC isoforms are detectable in glomeruli isolated from ZSF1 lean and obese animals, with no statistical difference between lean and obese for any PKC family member at the level of mRNA expression (**Figure 2A, and data not shown**).

**Figure 2.**
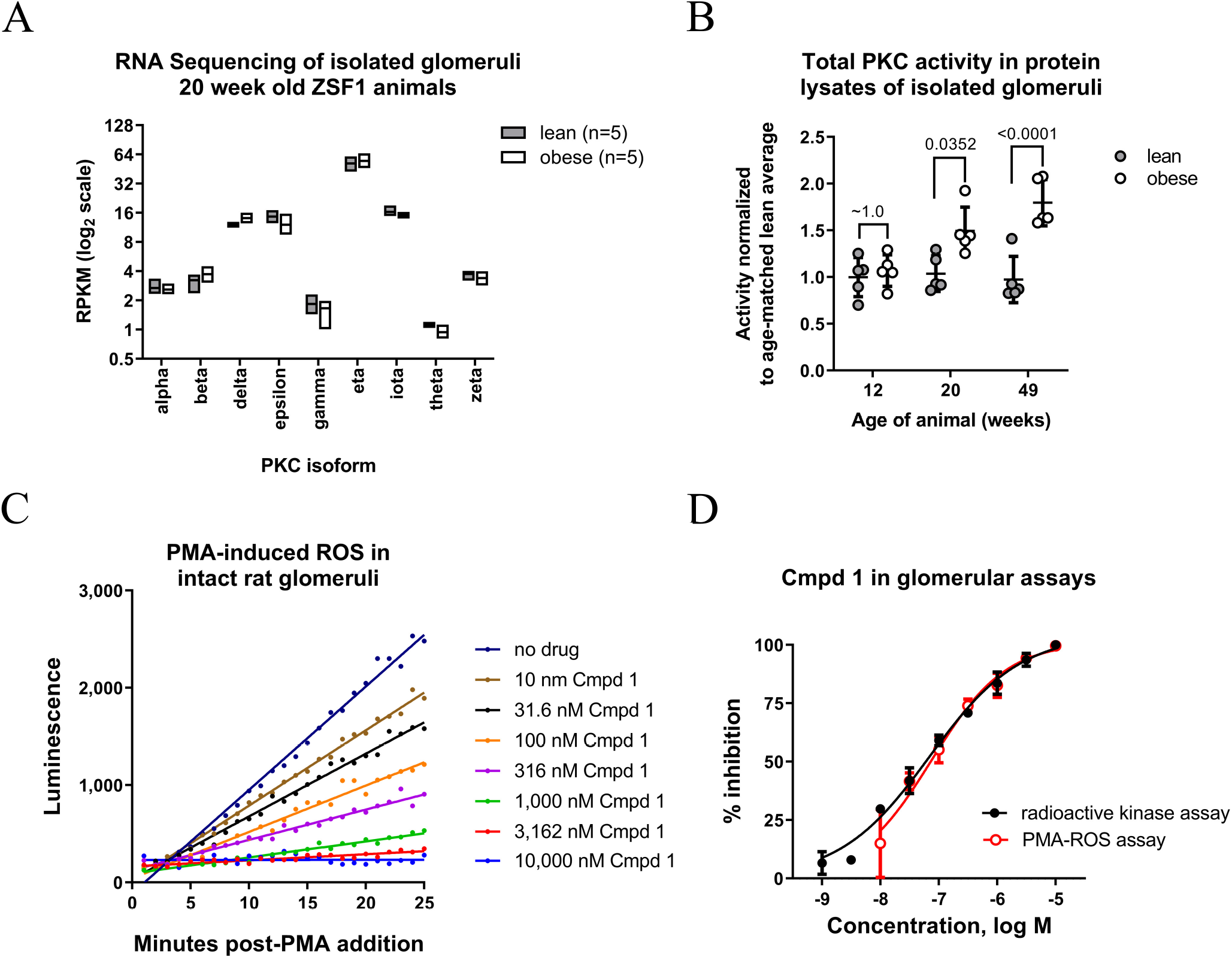
PKC activity in glomerular source material can be inhibited by MS-553 *in vitro*. (A) PKC mRNA isoform expression from RNA sequencing data of isolated glomeruli from lean and obese ZSF1 rats. The y-axis is reads per kilobase per million reads (RPKM), plotted in log2 scale. B) Pan-PKC activity in protein lysates made from glomeruli isolated from ZSF1 lean or obese rats at 12, 20, and 49 weeks of age. Activity was measured in an *in vitro* kinase assay using a pan-PKC pseudo-substrate peptide. C) Luminol assay for ROS production following stimulation of glomeruli isolated from SD rats with PMA, in the absence or presence of increasing amounts of Cmpd 1. D) Inhibition profiles of Cmpll in the radioactive kinase assay with glomerular lysates from 20-week-old ZSF1 obese animals, and in the PMA-ROS assay using intact glomeruli isolated from SD rats. Percent inhibition is normalized to no inhibitor (0%) versus 10 μM Cmpd 1 (100%). IC_50_ values for Cmpd 1 in these assays were 70 and 76 ⊓M, respectively.

To measure PKC activity and evaluate Cmpd 1 pharmacology in disease-relevant tissue, we performed *in vitro* assays using as glomeruli isolated by serially sieving minced kidney cortex as source material. In the first assay, protein lysates were generated from glomeruli isolated from ZSF1 lean or obese animals at 12, 20, and 49 weeks of age and used in a radioactive kinase assay with a pan-PKC pseudo-substrate peptide. By this analysis, PKC activity was similar in ZSF1 lean and ZSF1 obese animals at 12 weeks of age but became elevated in obese animals at 20 and 49 weeks of age (**Figure 2B**). In a second assay, glomeruli from Sprague Dawley (SD) rats were isolated, left intact, and then treated with the PKC activator PMA. PKC activity was then monitored by measuring reactive oxygens species (ROS) activity in real time by Luminol luminescence^14,56^ (**Figure 2C**). By performing the assay in the presence of increasing concentrations of Cmpd 1 and calculating the velocity of PMA-induced ROS at each compound concentration, an IC_50_ value of 76 nM was determined (**Figure 2D**). Similarly, when titrated into the aforementioned radioactive kinase assay using lysates from 20-week old ZSF1 obese glomeruli, Cmpd 1 had an IC_50_ value of 70 nM (**Figure 2D**). Although a limitation of both assays is that they do not distinguish which PKC family member is responsible for the measured response, the results indicate that Cmpd 1 can inhibit glomerular PKC activity that is intrinsic (radioactive kinase assay) or induced (PMA induced ROS), and does so with double-digit nM potency.

### Cmpd 1 administration reduces hyperphagia and body weights of ZSF1 obese rats

In preparation for a long-term efficacy study, and to facilitate dosing, we evaluated Cmpd 1 administration to ZSF1 obese animals through drug formulation in chow. Initial pharmacokinetic (PK) studies indicated that a chow formulation to deliver approximately 50 milligrams of Cmpd 1 per kilogram body weight (mpk) per day yielded suitable exposures (50 mpk/d; 0.744 g Cmpd 1 per kg chow; data not shown). We therefore characterized this chow dosing regimen in more detail. The results from a 4-day PK study with ZSF1 obese rats are shown in **Figure 3A**. Free drug concentrations in serum were roughly 700 nM and 400 nM in the morning and evening, respectively, consistent with higher feeding activity at night. Internal modeling indicated that these exposure levels approached or exceeded IC90 for both PKCα and PKCβ throughout the day (data not shown), and that this chow dosing regimen was suitable to test the intended mechanism.

**Figure 3.**
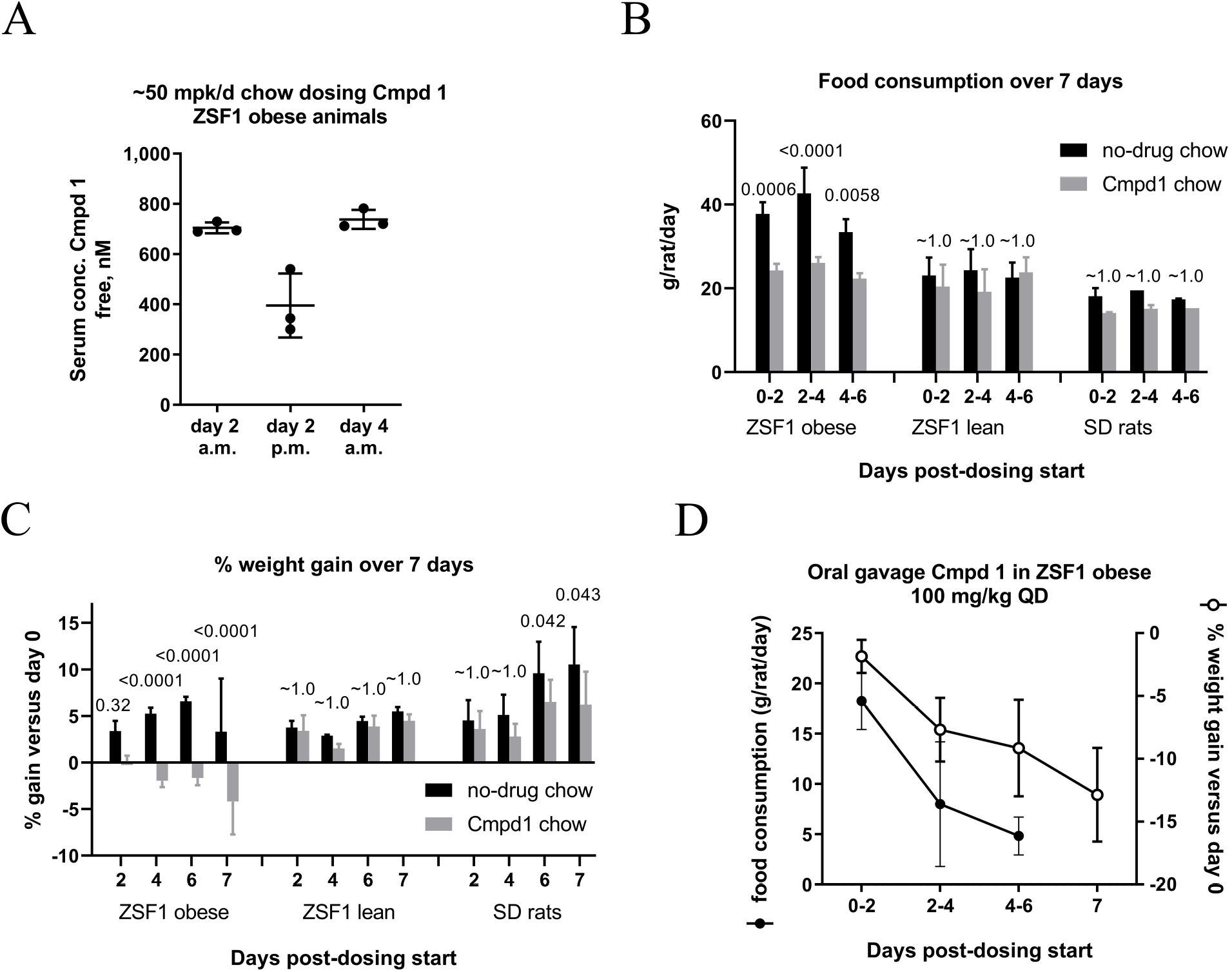
Pharmacokinetic (PK) measurement of Cmpd 1 when formulated in chow, and evidence for a food consumption and body weight effect with Cmpd 1 administration. A) Results from 4 day PK study for serum drug concentrations in the morning (9 a.m.) and late afternoon (4 p.m.) with chow dosing of Cmpd 1 to deliver an estimated 50 mpk/d. Food consumption (estimated; B) and % weight gain (C) over a 7 day study with ZSF1 obese, ZSF1 lean, or SD rats fed either no-drug chow or 50 mpk/d Cmpd 1 chow (3 animals for each ZSF1 group, 2 animals for each SD group), p-values for no-drug chow versus Cmpd 1 chow for each group are provided. (D) Average food consumption of no-drug chow (left y-axis; closed circles) and % weight gain (right y-axis; open circles) of ZSF1 obese rats with daily administration of 100 mpk/d Cmpd 1 by oral gavage (5 animals). Weights were measured on days 2,4, 6, and 7.

During these studies, we observed that ZSF1 obese animals appeared to consume less chow when it was formulated with Cmpd 1. To investigate this, we carried out a 7-day study in which we measured food consumption and weight gain by ZSF1 obese rats provided Cmpd 1 chow or regular (no-drug) chow. ZSF1 lean rats and SD rats were included for comparison. Over a 7-day study period, ZSF1 obese rats consumed less Cmpd 1 chow than no-drug chow based on periodic measurements of food provided versus food remaining in the hopper (**Figure 3B)**. ZSF1 obese rats consumed an average of 24 g/day Cmpd 1 chow versus 38 g/day no-drug chow. By comparison, ZSF1 lean and SD rats consumed an average of 23 g/day and 18 g/day of no-drug chow, respectively. Therefore, despite a 37% reduction in food intake by ZSF1 obese rats, the amount of Cmpd 1 chow consumed did not drop below normophagic levels. This effect of reduced food intake with Cmpd 1 chow was considerably more pronounced in ZSF1 obese rats than in ZSF1 lean or SD rats, although a trend towards reduced food intake was evident in these latter groups (**Figure 3B;** black versus grey bars). ZSF1 obese rats fed Cmpd 1 chow also lost weight over the 7-day study period in a manner that was absent or muted in ZSF1 lean and SD rats fed the same chow (**Figure 3C**).

To eliminate the possibility that these observations with ZSF1 obese rats were non-pharmacology related, for example due to a potentially unpleasant taste of Cmpd 1, we examined food consumption and body weights of ZSF1 obese animals administered Cmpd 1 by oral gavage (100 mpk daily; **Figure 3D**). Over 7 days, both food consumption and animal body weights decreased. Therefore, in 7-day studies with dosing through either chow or oral gavage, Cmpd 1 blunted the hyperphagic response and reduced the body weights of ZSF1 obese rats. Moreover, in this duration, the effects on food intake and body weight appeared to be considerably more pronounced in ZSF1 obese rats than in ZSF1 lean or SD rats.

### Design of 10-week efficacy study

Based on these findings and previous experience with the model, we designed a therapeutic efficacy study to evaluate chow dosing of Cmpd 1 for ten weeks initiating in 22-week-old ZSF1 obese rats. We also included a no-drug comparator group of ZSF1 obese rats for which food intake was restricted to approximately match that of Cmpd 1-dosed animals. To achieve this, the estimated amount of Cmpd 1 chow consumed each week was monitored and this amount of no-drug chow was placed in the hopper the following week for the comparator group. As a result, the estimated food intake amounts were staggered between these two groups of ZSF1 obese rats by one week. A summary of the study groups is provided in **Figure 4A**. Five age matched ZSF1 lean rats were included as a control (*Lean group*). The three groups of ZSF1 obese rats comprising the rest of the study consisted of the following: a group with no drug administration (*Obese group*), a group provided Cmpd 1 chow (*Cmpd 1 group*), and a group with estimated weekly food intake to match that of the Cmpd 1 group in the previous week (*Food matched group*). These three groups consisted of 11 or 12 animals at study start, three of which were sacrificed at study mid-point for interim analyses (data not shown). Food consumption and body weight were monitored weekly and non-fasting blood was taken for analysis at 0, 2.5, 5, 7.5, and 10 weeks. Animals were placed in metabolic cages at 0, 5, and 10 weeks for renal function tests and sacrificed after the 10-week study duration, corresponding to 32 weeks of age, for terminal histological and molecular analyses. **S**ubsequent analyses include all data for all available animals unless otherwise noted.

**Figure 4.**
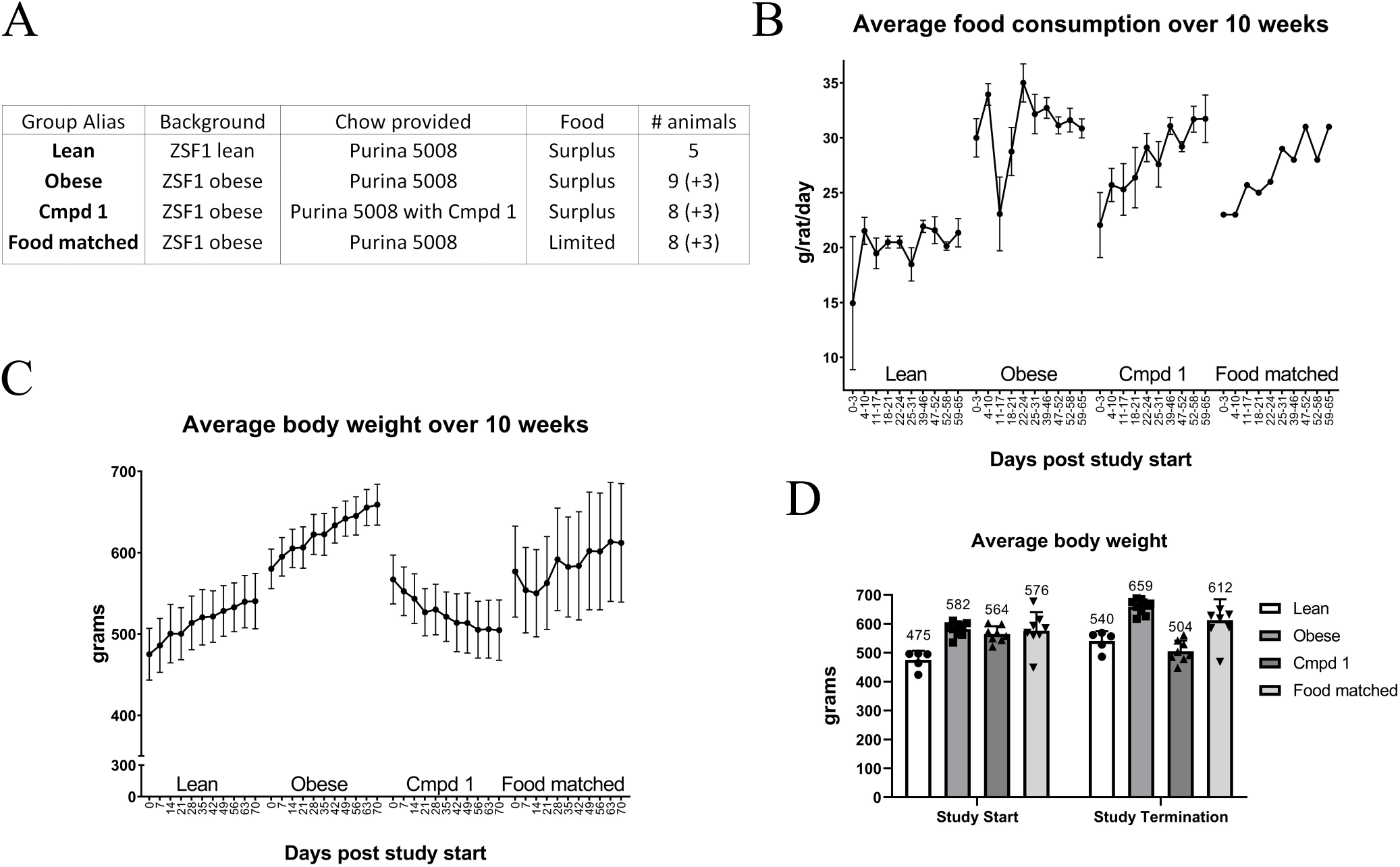
Study groups, food consumption, and body weight over the course of a 10 week efficacy study. (A) Study design. All animals were provided Pruina 5008 chow (27% kcal protein, 17% kcal fat, 57% kcal carboydrates) and enrolled at 21-22 weeks of age. The Cmpd 1 group was provided chow that was formulated with O.744g Cmpd 1 per kg chow (estimated 50 mpk/d). The amount of food provided to the Food matched group each week was determined by the estimated food consumption by the Cmpd 1 group the previous week. For all three ZSF1 obese groups, 3 animals were taken down at study mid-point for an interim analysis. (B) Average food consumption for all animals over the 10-week study duration as estimated by amount of food provided and remaining in the hopper for the indicated intervals. (C) Average body weight over the 10-week study duration for all animals. (D) Average body weights at study start and study termination (interim animals excluded), with average weights indicated numerically above the bars.

### Food consumption and animal body weights over study duration

As anticipated, Cmpd 1 animals displayed an altered food intake pattern. In the first three days, the estimated average food consumption by this group was 22 g/day, down from 30 g/day in the Obese group (27% reduction; **Figure 4B**). By comparison, the Lean group consumed an average of 15 g/day in this period. Food consumption by Cmpd 1 animals gradually normalized over the 10-week study duration, and by study termination these animals consumed an estimated 31 g/day compared to 32 g/day by the Obese group. Despite this normalization, and in clear contrast to the Food matched group, Cmpd 1 animals steadily lost weight over the entire study duration (**Figure 4C).** Indeed, at study termination, Cmpd 1 animals were on average approximately 7% less heavy and not statistically different than Lean animals (504 -/+ 35 grams, versus 540 -/+ 30 grams, p = 0.84; **Figure 4D**). Thus, the effect of Cmpd 1 in suppressing the hyperphagia of ZSF1 obese rats was also observed at the outset of this study but dissipated over time, presumably due to a compensatory increase in food intake as the animals lost weight.

### Non-fasting serum parameters

We profiled animals for non-fasting serum chemistry relevant to the T2D condition. Non-fasting conditions were used to minimize the impact of feeding disruptions on study outcome, and are a limitation of these measurements. ZSF1 obese rats displayed multiple features of metabolic syndrome at their study start age of 22 weeks. These included elevated total and LDL cholesterol, reduced HDL cholesterol, elevated triglycerides, elevated glucose, and elevated insulin (**Figure 5**). Both Cmpd 1 and Food matched groups displayed trends towards reduced LDL cholesterol and triglyceride levels, whereas little to no effects on serum total cholesterol, HDL cholesterol, and glucose were observed for these two groups. Although for most measurements there was no discernable difference between these two groups under these experimental conditions, a notable difference was observed in insulin levels. While Food matched animals displayed persistent hyperinsulinemia, Cmpd 1 animals displayed reduced insulin levels over much of the study duration. This reduction was prominent at weeks 2.5, 5, and 7.5 weeks but was less evident at 10 weeks, for reasons that are presently unclear. None-the-less, by example, at week 5 study midpoint, average insulin measurements for the various groups were as follows: Lean 0.26 (-/+ 0.16); Obese 9.3 (-/+ 3.5); Cmpd 1 0.62 (-/+ 0.35), and Food matched 9.9 (-/+ 6.2) ng/mL.

**Figure 5.**
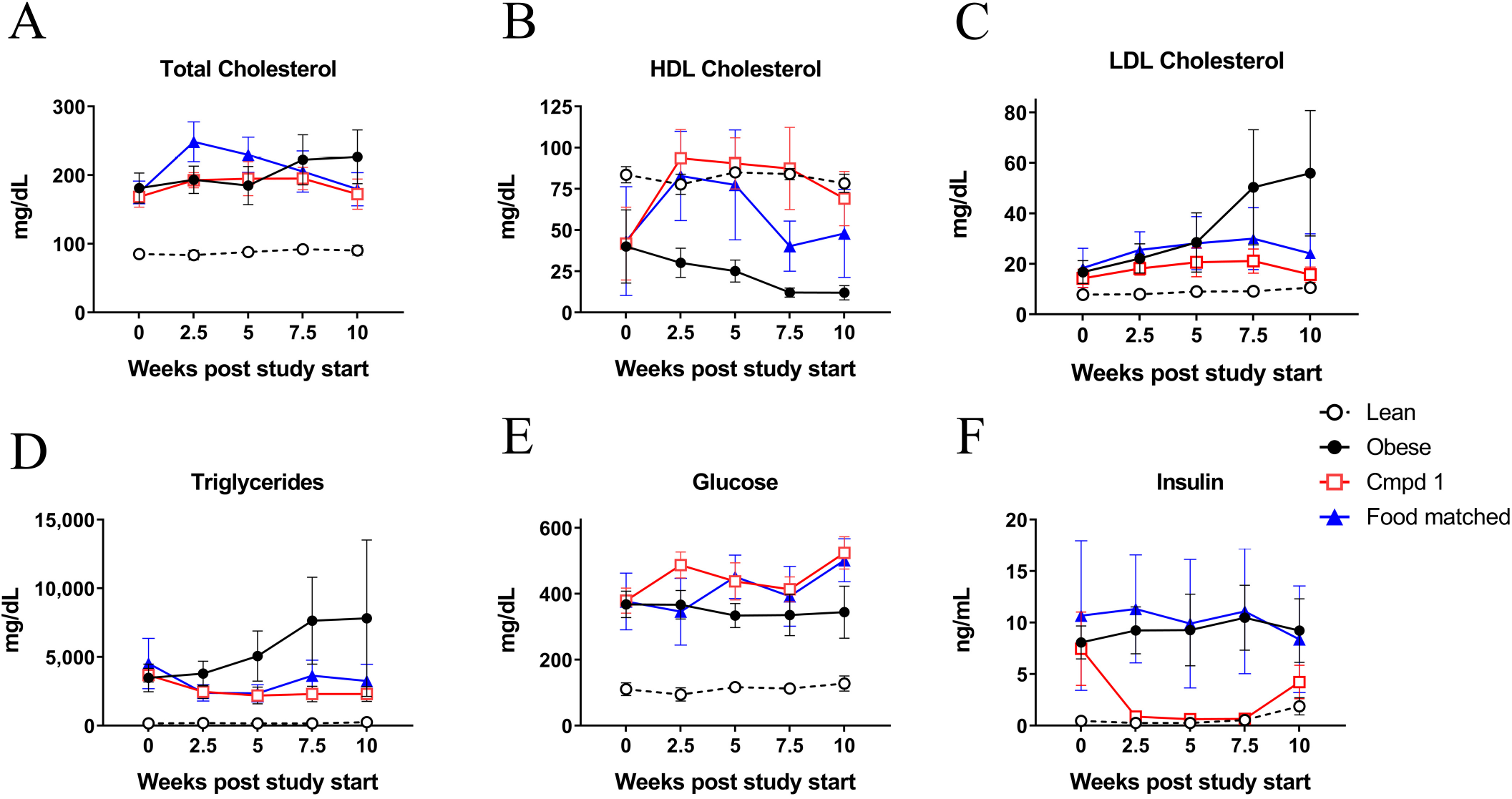
Non-fasting serum measurements at 0, 2.5, 5, and 10 weeks study duration. (A) Serum total cholesterol, (B) HDL cholesterol, (C) LDL cholesterol, (D) triglycerides, (E) glucose, and (F) insulin levels. Non-fasting conditions were used to minimize disruptions to feeding behavior during the study.

### Cmpd 1 halts DN progression indirectly through its effect on food intake patterns

Obese rats displayed renal impairment at their study start age of 22 weeks with approximately 300-fold greater microalbumin to creatinine ratio (uMALB:creatinine ratio) than Lean animals (14,520 -/+ 4,866 μg/mg, vs 45 -/+42 μg/mg; **Figure 6A**). Renal function in the Obese group progressively declined over the 10-week study period, as evidenced by steadily worsening uMALB:creatinine ratio and 24-hour urinary protein excretion (**Figures 6A, B**). In Cmpd 1 and Food matched groups, this progressive decline was virtually halted. Although the profile appeared slightly better in Cmpd 1 animals, we note that reduced food intake for the Food matched group was shifted by one week relative to Cmpd 1 animals over the study duration.

**Figure 6.**
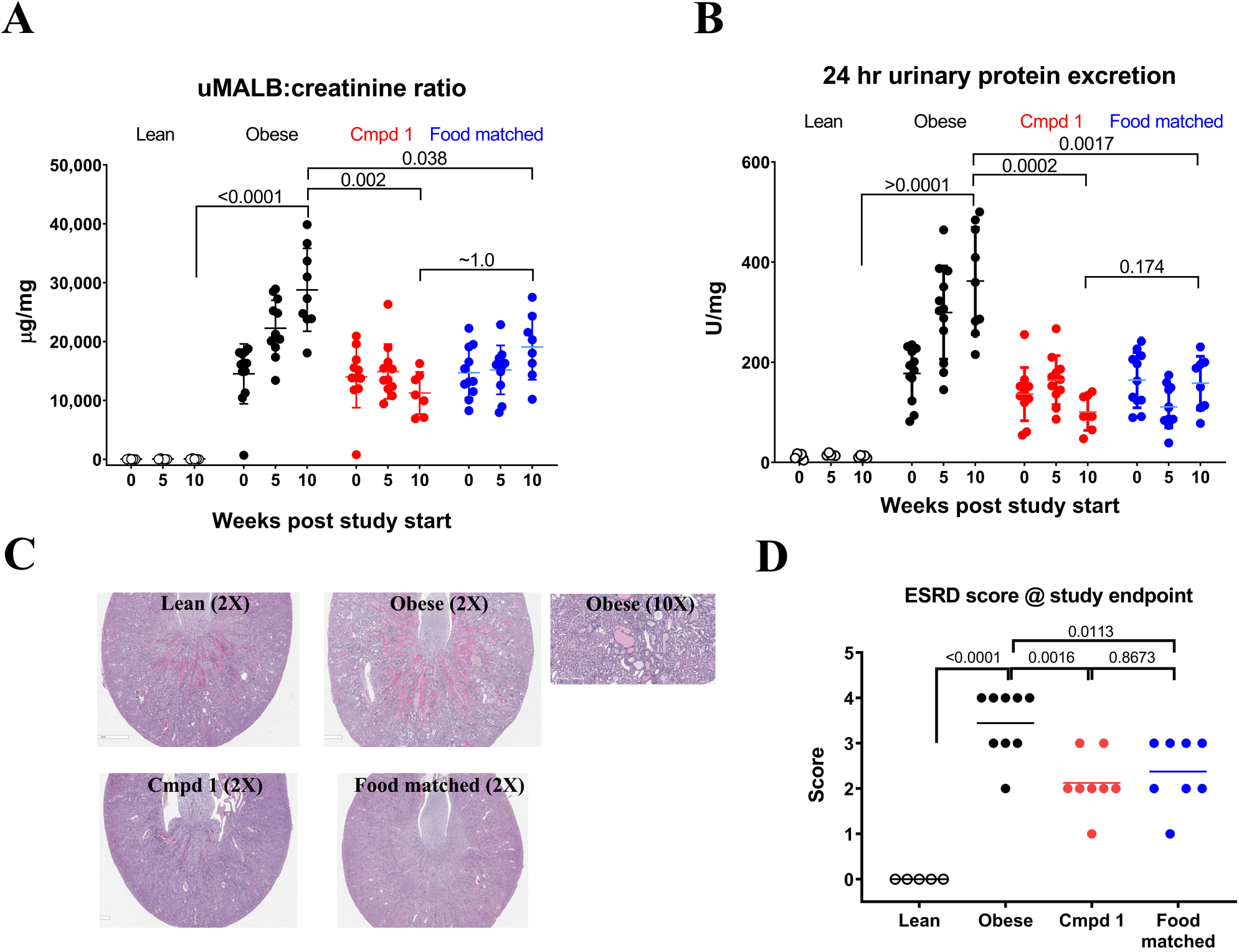
Renal endpoints over study duration. (A) Urinary microalbumin (uMALB) to creatinine ratio and (B) 24 hr urinary protein excretion at study start, midpoint, and endpoint. For clarity, relevant statistical analyses are provided only for the 10 week time-point in (A) and (B). We note that data are missing for one animal in the Cmpd 1 group at 10 weeks due to lack of urination by this animal in the metabolic cage used for these analyses. (C) Representative images of microscopic changes in the kidneys at study endpoint. Images were taken at 2X magnification; a representative 10X magnification of an obese animal is shown to highlight renal abnormalities. (D) ESRD score (severity grade 1-5; see text for details) following blinded analysis of the glomerular, tubular, and interstitial histological findings.

By histology, microscopic findings at study termination in the kidneys of Obese animals included changes in the glomeruli and tubulointerstitial compartments (**Figure 6C**). In the glomeruli, global to segmental multifocal mesangial expansion was observed with variable infiltrate of inflammatory cells, with and without synechiation, and thickening of the Bowman’s capsule. Tubular changes included hyaline protein casts within mostly medullary and cortico-medullary tubules, basophilic cortical and medullary tubules, tubular dilatation, attenuated basophilic epithelial cells, basement membrane thickening, occasional tubular hypertrophy within the cortex, and rare scattered mineralization. Variable expansion of the interstitium could also be observed, with mixed inflammatory cells and loose fibrous connective tissue. In a blinded analysis, the severity of kidney pathology (ESRD score) was graded from 1 (minimal) to 5 (severe; **Figure 6D**). ESRD scores at study termination were reduced comparably in both Cmpd 1 and Food matched groups, consistent with the urine biomarker analyses. We conclude that therapeutic dosing of Cmpd 1 halts but does not reverse renal function decline and does so indirectly, through its effects on the feeding behavior of ZSF1 obese rats.

### Cmpd 1 directly affects adiposity of liver and iWAT in ZSF1 obese rats

We examined two other tissues relevant to the model and our observations: liver and inguinal white adipose tissue (iWAT). Cmpd 1 animals had reduced liver weights; however, this reduction appeared to simply reflect differences in overall body weights (BW; liver to BW ratio; **Figure 7A**). Histological analyses, however, revealed differences in the livers across the groups. Obese animals had fatty livers, as exemplified in the representative histology images in **Figure 7B**. Microscopic findings included vacuolation, presumably from lipid accumulation, characterized as centrilobular to midzonal, and diffuse or multifocal macrovacuolation of hepatocytes with eccentric nuclei (consistent with lipid). To compare treatment groups, the severity scores of liver macrovacuolation were assigned in blinded fashion and ranged from 0 (no macrovacuolation) to 3 (moderate macrovacuolation; **Figure 7C**). Cmpd 1 reduced liver macrovacuolation scores in a manner that was not observed in the Food matched group: after 10-weeks of dosing, 5 of 8 Cmpd 1 animals had liver macrovacuolation scores of zero, as opposed to 1 of 8 animals in the Food matched group.

**Figure 7.**
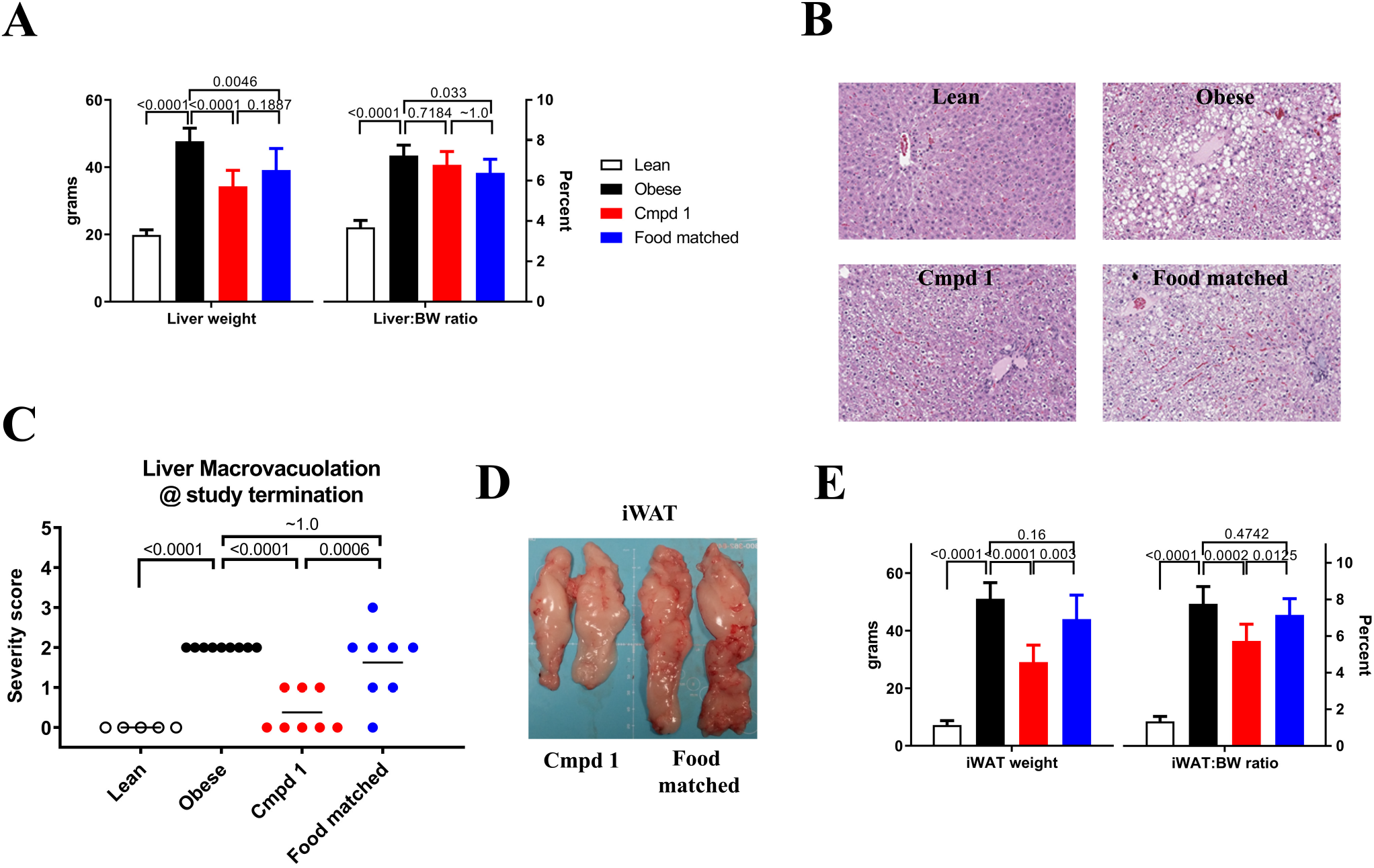
Cmpd 1 reduces liver macrovacuolation and iWAT size. (A) Liver weights and liver to terminal body weight ratio (Liver:BW ratio) at study termination. (B) Representative microscopic findings in the liver, including hepatocyte vacuolation, at study termination. (C) Liver macrovaculation score of all animals at study termination, on a scale of 1 to 3, obtained from blinded analysis. (D) Image depicting size difference in iWAT of a representative Cmpd 1 and Food matched animal. (E) inguinal white adipose tissue (iWAT) weights and iWAT:BW ratio at study termination. Color coding in panel (E) is the same as used in panel (A).

For iWAT, the size of the tissue was noticeably different across the groups (**Figure 7D**). Cmpd 1 animals had substantially lower iWAT weights (29 -/+ 5.5 g) than iWAT weights of Obese animals (51 -/+ 5.3 g) or Food matched animals (44 -/+ 7.8 g; **Figures 7D and E**), although still considerably higher than iWAT weights of Lean animals (7.23 -/+ 1.32 g). In contrast to the differences in liver weights, differences in iWAT weights were not entirely attributed to differences in overall body weights (iWAT to BW ratio; **Figure 7E**). Thus, Cmpd 1 reduced liver vacuolation and iWAT weights directly and independently of its effects on food intake, and in accordance with the body weight profiles observed over the study duration. Given the clear effect of Cmpd 1 on iWAT, this tissue was selected for more detailed analyses.

### Molecular and cellular changes in iWAT with Cmpd 1 administration

To gain mechanistic insights into the effects of Cmpd 1 on iWAT, we performed RNA sequencing on five randomly selected animals from each of the four groups. Differential expression analysis was conducted for two contrasts: Obese versus Lean (disease-associated changes), and Cmpd 1 versus Food matched (Cmpd 1-associated changes that were not due to changes in food intake). By this analysis, we identified 2,523 disease-associated differentially expressed genes (DEGs), 970 Cmpd 1-associated DEGs, and 795 DEGs that were present in both contrasts (**Figure 8A**). Hierarchical clustering by Euclidean distance of these 795 shared DEGs differentiated between Lean and Obese, and between Cmpd 1 and Food matched (**Figure 8A**). A clear, albeit partial, recovery towards the Lean profile was observed in the Cmpd 1 group. Canonical Ingenuity Pathway Analysis (IPA) identified multiple pathways related to immune function as being elevated in Obese animals (Z-score greater than zero) and conversely inhibited in Cmpd 1 animals (Z-score less than zero; **Figure 8B**). Further analysis using IPA Diseases & Functions annotations identified “quantity of adipose tissue”, “subcutaneous fat”, and “white adipose tissue” as being elevated with disease and inhibited by Cmpd 1, whereas “uptake of monosaccharide” was reduced with disease and rescued by Cmpd 1 (**Figure 8C**). Causal network analysis identified several networks that were either disease-associated (VEGFA and HIPK2, both down) or Cmpd 1-associated (FABP4 and PRKCB aka PKCβ, both down; PPARGC1B, up), and only one network, CD44, that was reciprocally regulated in the two contrasts (**Figure 8C**). The expression of PKCβ and its causal network genes identified by this analysis (the β-adrenergic receptors ADRB1 and ADRB2), and of the cell surface glycoprotein CD44 and its causal network genes (IL1RN, LGALS3, and MMP12), are provided in **Figures 8D and 8E**. The expression of HAS2, which catalyzes the synthesis of the CD44 ligand hyaluronic acid, and SPP1 (aka Osteopontin), also a ligand of CD44, are also shown in Figure 8E.

**Figure 8.**
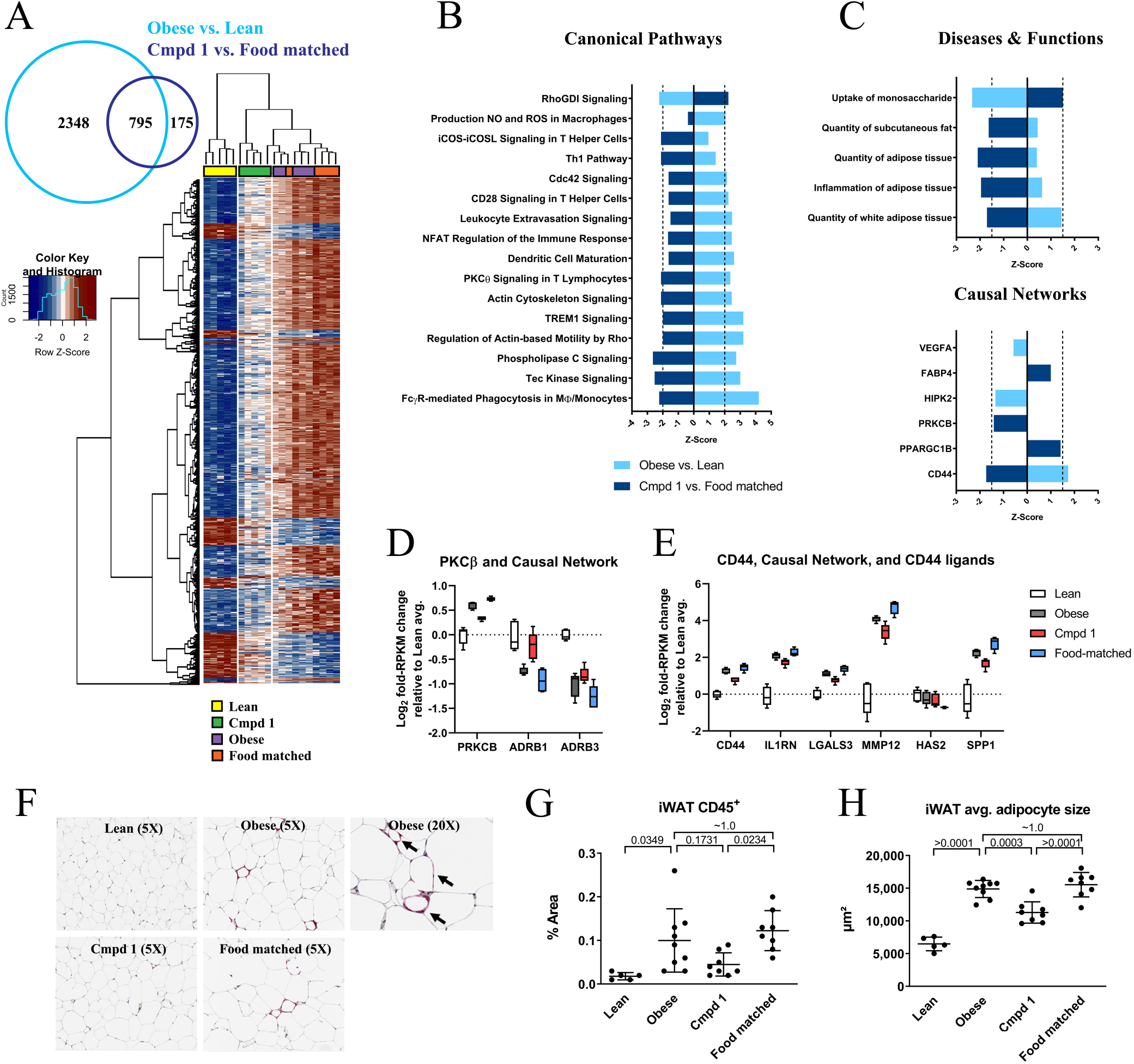
Transcriptomic and histological changes in iWAT with Cmpd 1 administration. (A) Venn diagram of differentially expressed genes (DEGs) in the two indicated contrasts. A heat map of unsupervised hierarchical clustering of the 795 DEGs present in both comparisons is shown. (B)Canonical pathways and (C) Diseases & Functions and Causal networks identified by Ingenuity Pathway Analysis (IPA) of these DEGs. Log2 fold-change relative to lean average of the indicated transcripts for two of the identified Causal Networks, PRKCB (aka PKCß; panel D) and CD44 (panel E). PKRCB was identified through changes in expression of ADRB1 and ADRB3. CD44 was identified through changes in expression of IL1RN, LGALS3, and MMP12. Two genes linked to CD44 ligands, HAS2 (hyaluron synthase 2) and SPP1 (osteopontin), are also shown. (F) Representative CD45 staining, in red, of iWAT tissue. A representative higher magnification image of an Obese animal is also provided, showing that CD45+ cells (arrows) resided adjacent to and encircling adipocytes. (G) Quantitation of CD45+ cells n whole slides using digital image analysis; data represents percent of iWAT area that is reactive on CD45 IHC. (H) Quantification of adipocyte size, in μm^2^, in HE-stained iWAT for all animals.

A clear finding from pathway analysis was a reduction in inflammatory signature in iWAT upon Cmpd 1 administration (Figure 8B), perhaps reflecting reduced inflammatory cell numbers. To evaluate this, we performed immunohistochemistry using the pan-leukocyte marker CD45. CD45-positive (CD45^+^) cell numbers in iWAT increased with disease, with positive cells tending to reside adjacent to and encircling adipocytes (**Figure 8F**). Average CD45^+^ cell staining area was lower in Cmpd 1 animals relative to Food Matched animals (0.045 -/+ 0.025 percent, versus 0.122 -/+ 0.04 percent); **Figure 8G**). Although variability within the Obese group precluded clear comparisons, an overall trend towards fewer CD45^+^ cells in iWAT was evident in the Cmpd 1 group. This suggests that the effects on inflammatory pathways observed by molecular profiling were due, at least in part, to reduced inflammatory cell numbers in iWAT with Cmpd 1 administration. In addition, the histological analysis revealed that adipocytes assumed a morphology consistent with white fat and were increased in size in the Obese group, and that these effects were blunted in the Cmpd 1 group but not in the Food matched group (**Figure 8H**). In summary, consistent with its macroscopic effect on the mass of iWAT tissue, Cmpd 1 had direct effects on inflammatory status and adipocyte morphology in this tissue in ZSF1 obese rats.

## Discussion

Here we describe what is, to our knowledge, the first report of therapeutic dosing of a dual small molecule inhibitor of PKCα and PKCβ in a rodent T2D model that is characterized by renal impairment that progresses to ESRD. In pilot studies, we observed that Cmpd 1 blunted the hyperphagia response of ZSF1 obese rats. This led to the inclusion of a comparator group in the 10-week efficacy study for which food consumption was restricted to approximate that of Cmpd 1-administered animals. Although food consumption of Cmpd 1-administered animals normalized over the study duration, this altered food intake pattern resulted in clear protection from renal function decline. In contrast to this marked but indirect effect, Cmpd 1 had direct effects on body weight, non-fasting serum insulin levels, liver macrovacuolation, and iWAT parameters that were not attributed to altered food consumption patterns.

We do not presently understand the striking effect of Cmpd 1 on the hyperphagia response of ZSF1 obese rats. Animals such as ZSF1 obese rats that are deficient in leptin receptor signaling are hyperphagic likely due, at least in part, to an inability to decrease AMPK activity in the hypothalamus, a normal consequence of the satiety hormone leptin^57,58^. PKCs have been implicated in inhibiting AMPK, in particular when DAG levels are elevated such as in conditions of hyperglycemia and hyperlipidemia^59^. Inhibition of hypothalamic PKCs, therefore, could mimic leptin action. Although based on its structural features Cmpd 1 is not expected to pass the blood-brain barrier, we cannot exclude an effect of Cmpd 1 in the hypothalamus. In addition, potential changes to the blood-brain barrier under conditions of metabolic syndrome would need to be taken into consideration^60^.

PKCα, for which research has focused more on cardiovascular dysfunction, immune-mediated arterial thrombosis, and cancer, has not been linked to body weight or adipose tissue homeostasis in the literature^61^. However, previous work using PKCβ deficient mice, both in a wild type background and in the background of leptin deficiency (ob/ob mice), revealed a clear role for PKCβ in adiposity and obesity^62–65^. These effects were attributed to an increase in fatty acid oxidation and altered balance between energy consumption and energy expenditure with PKCβ deficiency. In both wild type and ob/ob backgrounds, deletion of PKCβ led to reduced adiposity, decreased hepatic steatosis, and decreased visceral and iWAT mass without concomitant reduction in food consumption. While this is in apparent contrast the effects on hyperphagia we observe in our pharmacologic study, we note that food intake by Cmpd 1 animals, while initially depressed, recovered to normal levels by the 10 week time-point. No data on renal endpoints were presented in the ob/ob mouse background, a T2D model that displays features of early-to-moderate DN^66^. None-the-less, we have pharmacologically recapitulated the genetic observations with mouse PKCβ deficiency that link this PKC to body mass, fatty acid oxidation, fat storage, and adipose tissue remodeling. In addition, pharmacologically, Ruboxistaurin caused weight loss in (mRen-2)27 transgenic hypertensive rats but only when co-administered STZ, suggesting that the weight loss effect was linked to the T1D condition^67^. However, in another report, Ruboxistaurin prevented weight gain in animals administered the anti-psychotic drug clozapine, and did so without impacting the slightly elevated food consumption patterns with clozapine administration^68^. To our knowledge, clinical trials with Ruboxistaurin were not associated with weight loss or reduced adiposity in humans; however, our internal modeling indicates that Ruboxistaurin drug concentrations in these trials may not have afforded the levels of target suppression that we are likely achieving in our animal studies (data not shown). Given the striking effect of Cmpd 1 on hyperphagia in the present study, and of Cmpd 1 and PKC inhibitors on body weight and adiposity in multiple pre-clinical contexts, future work should evaluate whether these phenomena are model- or rodent-specific or more general in nature.

From RNA sequencing of iWAT we identified CD44 as a potentially important causal disease network that is induced by disease and reversed by Cmpd 1. CD44 is a cell-surface receptor for glycoproteins such as hyaluronan and SPP1 (osteopontin) and has a role in leukocyte migration and activation. Interestingly, in a high fat diet mouse model, CD44 blockade with a neutralizing antibody prevented obesity, reduced insulin resistance, and ameliorated adipose tissue inflammation^69^. While a link between PKC and CD44 is not conclusively known, it has been reported that activated PKCs phosphorylate CD44 to enable intracellular interactions between the CD44 cytosolic domain and the cytoskeleton, which is critical for CD44-induced cell motility^70^. Therefore, PKC inhibition could negatively impact CD44 action, resulting in reduced leukocyte accumulation. RNA sequencing revealed another network of potential interest, the β-adrenergic receptors ADRB1 and ADRB3, whose mRNA levels were downregulated with disease and rescued by Cmpd 1. This result is what implicated the PKCβ pathway in the causal network analysis. As ADRB1 and ADRB3 have been linked to promoting the proliferation of brown adipocytes and enhancing lipolysis^71,72^, this could constitute a potential mechanism for our findings with Cmpd 1 administration and PKC inhibition on iWAT size and adipocyte morphology, as has been proposed previously^63^.

Of interest to the current study are previous studies in humans using Ruboxistaurin, also known as LY-333531. Ruboxistaurin has been shown to improve renal outcome in multiple rodent models of diabetes^15,29,73,74^. In large clinical trials, Ruboxistaurin at 32 mg/d for up to 3 years in diabetic individuals showed some benefit in diabetic eye complications, for which the trials were designed, but did not improve kidney function in post-hoc analyses^75–77^. We note, however, that although Ruboxistaurin is touted to be a PKCβ-selective inhibitor, it is rapidly metabolized to the much less-selective desmethylruboxistaurin *in vivo*^78–80^. In-so-far as multi-PKC inhibition but not pan-PKC or multi-kinase inhibition may be desirable to prevent diabetes-induced tissue damage, studies with Ruboxistaurin are consequently difficult to interpret with respect to whether the touted mechanism was selectively or suitably tested. Alternatively, and as mentioned previously, our internal modeling indicates that Ruboxistaurin may not have achieved the drug concentrations and levels of target suppression in humans that we are achieving in our rodent studies.

We note several limitations of the current study. As with most studies using ATP-competitive inhibitors of kinases, off-target pharmacology is a concern. In addition, Cmpd 1 has activity against PKCθ, which has been linked to insulin sensitivity and obesity^81,82^. We attempted to minimize inhibition of kinases other than PKCα and PKCβ through chow dosing to avoid high peak-to-trough compound concentrations, but these limitations persist. Furthermore, we cannot assign observations from this study to PKCα inhibition, PKCβ inhibition, or inhibition of both kinases; indeed, as the literature strongly suggests that at least some of our observations can be linked to PKCβ inhibition, it is presently unclear to what extent PKCα inhibition is playing a role in any of our findings. A final and important limitation is that we were unable to evaluate a direct effect of Cmpd 1 administration on DN progression due to its effects on food consumption patterns in this model setting.

Beyond the effect on hyperphagia, and resulting protection from renal function decline, we demonstrate that Cmpd 1 has direct effect on body weights of ZSF1 obese animals and directionally beneficial effects on serum insulin, liver vacuolation, and disease-associated derangements in iWAT. Future studies should examine the precise mechanism underlying the effect of Cmpd 1 on these parameters, whether they translate to non-leptin models of obesity, and whether they can be reasonably expected to extend to humans in particular T2D individuals. In addition, although obese ZSF1 rats display liver steatosis, they do not progress to nonalcoholic fatty liver disease (NASH)^83^. Given the rising incidence of NASH, for which T2D is a significant risk factor, and the beneficial effects we observe with respect to adiposity and inflammatory cell infiltration in adipose tissue, evaluation of Cmpd 1 or related PKC inhibitors in a pre-clinical NASH setting may be warranted.

## Abbreviations

PKC: protein kinase C
DN: diabetic nephropathy
T2D: type 2 diabetes
T1D: type 1 diabetes
ESRD: end stage renal disease
STZ: streptozotocin
AGE: advanced glycosylation end products
ROS: reactive oxygen species
PMA: phorbol 12-myristate 13-acetate
PK: pharmacokinetic
IC_50_: half maximal inhibitory concentration
RPKM: reads per kilobase per million reads
iWAT: inguinal white adipose tissue
DEG: differentially expressed gene
IPA: Ingenuity Pathway Analysis

